# *Srsf2^P95H/+^* co-operates with loss of TET2 to promote myeloid bias and initiate a chronic myelomonocytic leukemia like disease in mice

**DOI:** 10.1101/2022.06.21.496931

**Authors:** Jane Jialu Xu, Alistair M. Chalk, Meaghan Wall, Wallace Y. Langdon, Monique F. Smeets, Carl R. Walkley

## Abstract

Recurrent mutations in two pathways - the RNA spliceosome (eg. *SRSF2, SF3B1, U2AF1*) and epigenetic regulators (eg. *DNMT3, TET2*) – contribute to the development of myelodysplastic syndrome (MDS) and related myeloid neoplasms. In chronic myelomonocytic leukemia (CMML), *SRSF2* mutations occur in ∼50% of patients and *TET2* mutations in ∼60%, representing two of the most frequent mutations in these cancers. Clonal analysis has indicated that either mutation can arise as the founder lesion, however, our understanding of the basis for the co-operativity of these mutations in the evolution of CMML is limited. Based on human cancer genetics we crossed an inducible *Srsf2*^*P95H/+*^ mutant model with *Tet2*^*fl/fl*^ mice to mutate both concomitantly (or individually) in hematopoietic stem cells. At 20-24 weeks post gene mutation, we observed subtle differences in the *Srsf2*/*Tet2* mutants compared to either single mutant. Under conditions of native hematopoiesis with aging, we see a distinct myeloid bias and monocytosis in the *Srsf2*/*Tet2* mutants. A subset of the compound *Srsf2*/*Tet2* mutants display an increased granulocytic and distinctive monocytic proliferation (myelo-monocytic hyperplasia), with increased immature promonocytes and monoblasts (∼10-15% total nucleated cells), and evidence of binucleate promonocytes. Exome analysis of progressed disease demonstrates mutations in genes and pathways similar to those reported in human CMML. Upon transplantation, recipients developed leukocytosis, monocytosis and splenomegaly. This demonstrates we can reproduce *Srsf2/Tet2* co-operativity *in vivo*, yielding a disease with core characteristics of CMML, unlike single *Srsf2* or *Tet2* mutation. This model represents a significant step toward building high fidelity and genetically tractable models of CMML.

**Key points:**

- *Srsf2*^P95H/+^ co-operates with *Tet2*^*-/-*^ to initiate CMML in a murine model
- *Srsf2*^*P95H*^ and *Tet2* null mutations synergize in the development of monocytosis

## Introduction

The genomic landscape of myelodysplastic syndromes (MDS) and related myeloid neoplasms is described. This reveals patterns of recurrent mutations within individual genes and across gene classes. While the roles of many of the identified genes have been studied individually in animal models, the basis for co-operativity is less well understood. Valuable insight can be gained from investigating these genetic co-operativities, especially between driver mutations. This will improve understanding of the antecedent clonal hematopoiesis and its evolution to myeloid biased neoplasia. It will also inform understanding of how the distinct functional pathways regulate hematopoiesis, allowing a more accurate modeling of the human disease and the development of high fidelity and high penetrance models of human cancer.

In MDS and MDS/MPN co-mutation of *SRSF2* and *TET2* are profoundly enriched (p=2.63×10^−14^; q=1.21×10^−12^; cohort of n=3047)^1-4^. In chronic myelomonocytic leukemia (CMML), *SRSF2* and *TET2* are the most frequently occurring mutations (50% and 60%, respectively) and are the likely founder mutations in a significant proportion of cases^5^. In CMML and the related chronic neutrophilic leukemia and atypical chronic myeloid leukemia, there is a positive association between mutation of *SRSF2* and *TET2* (p=0.002; n=218)^6^. This association is strongest in CMML (p=0.004; 52 co-mutated of 146 samples)^6^. Thus, establishing an *in vivo* model of these mutations could generate a valuable tool to study CMML initiation and evolution.

CMML is a rare cancer with an incidence of 3 cases per 100,000 individuals older than 60 years (0.5/100,000 p.a. in all ages)^4,7^. Median survival rates for CMML are between 24-97 months for low-risk forms, to 5-18 months for high-risk^8^. Approximately 15-20% of patients will undergo leukemic transformation and these patients have 5-year survival of <10%^9^. There is no current consensus treatment for CMML, and current therapy revolves around hydroxyurea or hypomethylating agents but clinical responses are generally not sustained. The development of preclinical models is needed to advance the understanding of CMML and to provide a platform for progressing testing of new therapeutic approaches.

Modeling CMML *in vivo* using human patient derived xenografts produces highly variable cell engraftment in immunocompromised murine recipients^10,11^. An alternative is to use genetically modified immunocompetent mice and engineer the relevant mutations, an approach already applied to several of the genes recurrently mutated in CMML^12-17^. These predominantly single gene mutation models partially deciphered the consequences of each lesion but have to date not recapitulated the clinical landscape of the multiple dysfunctional pathways occurring in CMML^18,19^. Yoshimi et al., demonstrated experimentally an interaction of either *SRSF2* or *TET2* mutation with tumor associated point mutations of *IDH2* with these genetic combinations also noted in *de novo* AML^10^. They did not report co-operativity between the *Srsf2* mutant mouse model used and a loss of function *Tet2* mutation, even though *SRSF2* and *TET2* mutations are highly correlated in human CMML^1,2,4,5^.

To explore the interaction of *Srsf2*^*P95H/+*^ and *Tet2* mutations, we generated a conditionally mutated *Srsf2*^*P95H/+*^ *Tet2*^*fl/fl*^ model. We used concurrent inducible somatic *Srsf2*^*P95H/+*^ mutation^17^ and deletion of *Tet2*^15^ within the hematopoietic stem cell populations to assess how these mutations impact hematopoiesis and enable development of myeloid neoplasia. We observed that a subset of double mutant mice developed increased granulocytes and monocytes (myelomonocytic hyperplasia) with elevated immature promonocytes and monoblasts (∼10 – 15% total nucleated cells in the bone marrow). These features are consistent with CMML, demonstrating that this model is a significant first step toward modelling CMML and for pre-clinical therapeutic testing.

## Methods

All animal experiments were approved by the Animal Ethics Committee, St. Vincent’s Hospital, Melbourne, Australia (AEC#001/16 and 007/19).

### Mice

The *Srsf2*^*P95H/+*^ (C57BL/6NTac-*Srsf2*^*tm2874(P95H)Arte*^), *Tet2*^*fl/fl*^ mice (B6;129S-*Tet2*^*tm1*.*1laai*/J^; strain #017573, Jackson Laboratory), *c-Cbl*^*-/-*^ (MGI:2180578)^20^, *Rosa26*-CreER^T2^ (strain #008463, Jackson Laboratory) and h*Scl*-CreER^T^ have been previously described^15,17,21^. h*Scl*-CreER^T Tg+^ *Srsf2*^*P95H/+*^ mice were crossed to *Tet2*^*fl/fl*^ mice to generate h*Scl*-CreER^T Tg+^ *Srsf2*^*P95H/+*^ *Tet2*^*fl/fl*^ mice. Congenic B6.SJL-Ptprc^aPep3b/BoyJArc^ were purchased form the Animal Resources Centre (Canning Vale, Western Australia). Heterozygous CD45.1/CD45.2, derived from the intercross of C57BL/6 and B6.SJL-Ptprc^aPep3b/BoyJArc^ mice, were bred at St. Vincent’s Hospital. Tamoxifen containing food was prepared containing 400mg/kg tamoxifen citrate (Sigma or Selleckchem) with 5% sugar (refined white sugar) in irradiated standard mouse chow (Specialty Feeds, Western Australia). Tamoxifen containing chow was fed *ad libitum* during the treatment period then animals were returned to normal diet.

### Tissue collection and Immunophenotyping

For monitoring of animals, peripheral blood (PB) samples were obtained via retro-orbital bleeding. For analysis at the end of the experiment or when animals were moribund, PB was collected, and mice then euthanized by CO_2_ asphyxiation or cervical dislocation. Bone marrow (BM) was flushed from both femurs. Spleen and thymus were weighed and crushed through 40µM cell strainers (BD). PB, bone marrow, spleen and thymus suspensions were counted on a Sysmex KX21 hematological analyzer (Sysmex Corp). For immunophenotyping, cells were analyzed on a BD LSRIIFortessa (BD Biosciences) and data was analyzed using FlowJo^17,22,23^.

### Bone marrow transplantation

All transplantations were performed into lethally irradiated (2 × 5Gy; 3 hours apart; Gammacell Irradiator) B6.SJL-Ptprc^aPep3b/BoyJArc^ recipients (Animal Resources Centre, Perth, Australia; 3-6 recipients per transplantation cohort) through intravenous injections. For competitive transplants, BM from heterozygous CD45.1/CD45.2 was used as competitor. Recipient animals received antibiotics (Enrofloxacin, Baytril) in the drinking water for 3 weeks after irradiation. Experiments were performed in duplicate or as indicated.

### Histology analysis

Blood films were prepared with 2µl of PB, air-dried and fixed in 100% methanol. For BM cytospins, 1×10^5^ whole BM cells in 100µL PBS were spun onto charged microscope slides in a Cytospin™4 Cytocentrifuge (Thermo Fisher), air-dried and fixed in 100% methanol. May-Grünwald-Giemsa staining was performed as described by the manufacturer. The slides were imaged on a Leica DMRB fluorescence microscope at 20x or 40x magnification and processed with cellSens software (Olympus).

### Exome sequencing

Exome sequencing was performed on genomic DNA from whole BM of 11 independent animals from 50 to 60 weeks post–tamoxifen treatment or when moribund. DNA was isolated using Gentra Puregene kit (Qiagen) as described by the manufacturer. Exome capture was performed by GENEWIZ with the Agilent SureSelect Mouse Exon system and sequenced on the Illumina platform generating 150bp PE reads (GENEWIZ Ltd, Hong Kong). Reads were mapped using BWA mem (v0.7.15-r1140) vs mm10 using default parameters^24^. Depth of coverage was calculated using mosdepth^25^. Variants were called using Strelka^26^, with input from structural variants called first using Mantra^27^. Variants were filtered using snpSift^28^and variant effects were annotated using snpEff (v3.4t)^29^.

### Statistical Analysis

The results were analyzed using the one-way ANOVA or unpaired Student t test unless otherwise stated using Prism or Microsoft Excel software. *P* <0.05 was considered significant. All data are presented as mean ± standard error of the mean.

### Data sharing statement

The exome sequence data are available at GEO under accession number: Raw data are available in the Sequence Read Archive (SRA) submission: PRJNA850841.

## Results

### *Srsf2^P95H/^*^+^ *Tet2*^-/-^ double mutant mice develop myeloid expansion and macrocytosis

To generate the *Srsf2*^*P95H/+*^ and *Tet2* compound mutant model, we crossed our characterized *Srsf2*^*P95H/+*^ model^17^ with a conditional *Tet2* loss of function allele (*Tet2*^*fl/fl*^)^15^ to generate h*Scl*-CreER^T^ *Srsf2*^*P95H/+*^ *Tet2*^*fl/fl*^ cohorts (Figure 1A). At 8-10 weeks of age, the mice were fed tamoxifen containing food for 4 weeks and then we performed a full hematological assessment at 20 weeks and at 56 weeks post gene mutation. All parameters are compared to age-matched tamoxifen-treated Cre-positive wild-type (WT) controls. At 20 weeks post gene mutation, there were macrocytic changes in the peripheral blood (PB) red blood cells (RBC; Supplemental Figure 1A-D). The hemoglobin level (Hb) was elevated in the double mutant, while the RBC numbers dropped in the *Srsf2*^*P95H/+*^ single mutant mice (Supplemental Figure 1B). There was a decreased bone marrow (BM) cellularity in both the double mutant and *Srsf2*^*P95H/+*^ single mutant mice (Supplemental Figure 1E). There were no other changes in the PB or BM parameters at 20 weeks post gene mutation (Supplemental Figure 1B-G).

**Figure 1.**
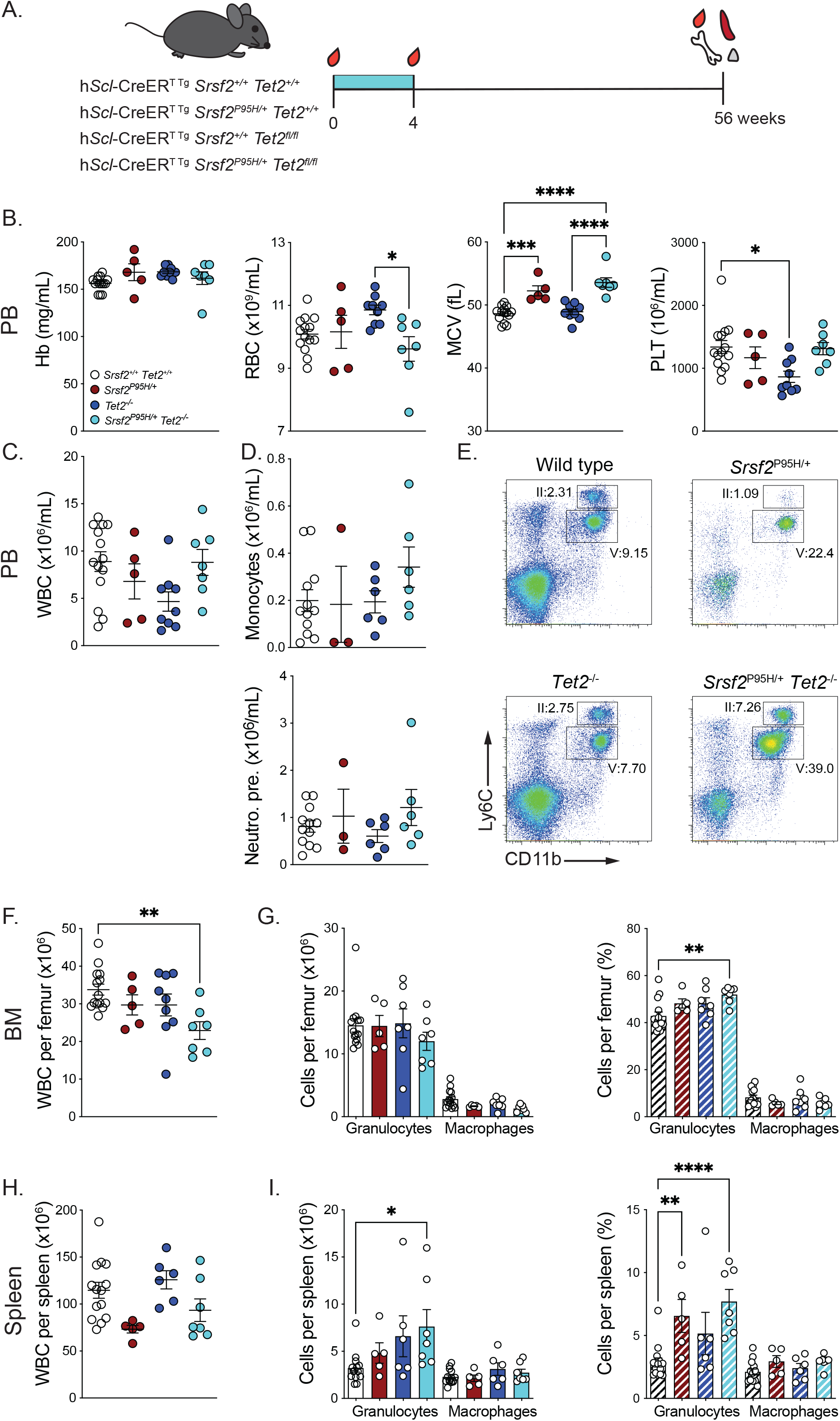
Concurrent mutation of *Srsf2*^*P95H/+*^ and deletion of *Tet2* leads to myeloid bias in the bone marrow and spleen. (**A**) Schematic of h*Scl*-CreER *Srsf2*^P95H/+^ *Tet2*^-/-^ experiments. The blue box indicates the tamoxifen administration period. (**B**) PB indices after 52 weeks of Cre activation. (**C**) Peripheral white blood cell count. (**D**) Number of monocytes (Ly6C^hi^ CD11b+) and neutrophil precursors (Ly6C^med^ CD11b+) in the peripheral blood. (**E**) Representative FACS plot of monocytes (II) and neutrophil precursors (V) in the peripheral blood of each genotype. (**F**) *Srsf2*^P95H/+^ *Tet2*^-/-^ bone marrow cellularity compared to age matched control. (**G**) Number and percentage of granulocytes and macrophages in the bone marrow. (**H**) *Srsf2*^P95H/+^ *Tet2*^-/-^ spleen cellularity compared to age matched control. (**I**) Number and percentage of granulocytes in the spleen of *Srsf2*^P95H/+^ *Tet2*^-/-^ mice. N=5-13 per genotype. Presented as mean +/-standard error of mean. One-way ANOVA performed against wild-type cells. \**P*<.05, \*\**P*<.01, \*\*\**P*<.001, \*\*\*\**P*<.0001.

At 56 weeks post tamoxifen, the PB of the double mutant cohort showed persistent macrocytosis and a trend toward an increased number of monocytes, even though the total leukocyte number remained unchanged (Figure 1B-E). There was hypocellularity of the BM of the double mutant mice (Figure 1F). No changes were observed in the absolute number of myeloid cells per femur, but the *Srsf2/Tet2* compound mutant mice had an increased percentage of granulocytes (Figure 1G). While the total cellularity of the spleen remained unchanged, the double mutant mice had increased granulocytes, indicative of myeloid expansion in the spleen (Figure 1H-I). A similar trend was observed in the *Srsf2*^*P95H/+*^ and *Tet2*^*Δ/Δ*^ single mutant mice, except that these genotypes exhibited a normal BM cellularity with an elevated percentage of granulocytes in the spleen. In contrast, there was extensive lymphoid suppression from the pre-B cell stage to mature B cells with an increasing severity from the single to the double mutants. Proportionally there was an expansion of the pre-pro B cells in the double mutant BM while the pre-B cells decreased, suggesting an impairment in pre-pro B cell differentiation (Figure 2A). Similarly, the nucleated erythroid progenitors (Ter119^+^CD71^hi^) and late-stage erythroid cells (Ter119^+^CD71^low^) were decreased in the compound mutant mice (Figure 2B). The T lymphoid populations remained unchanged in all genotypes (Data not shown).

**Figure 2.**
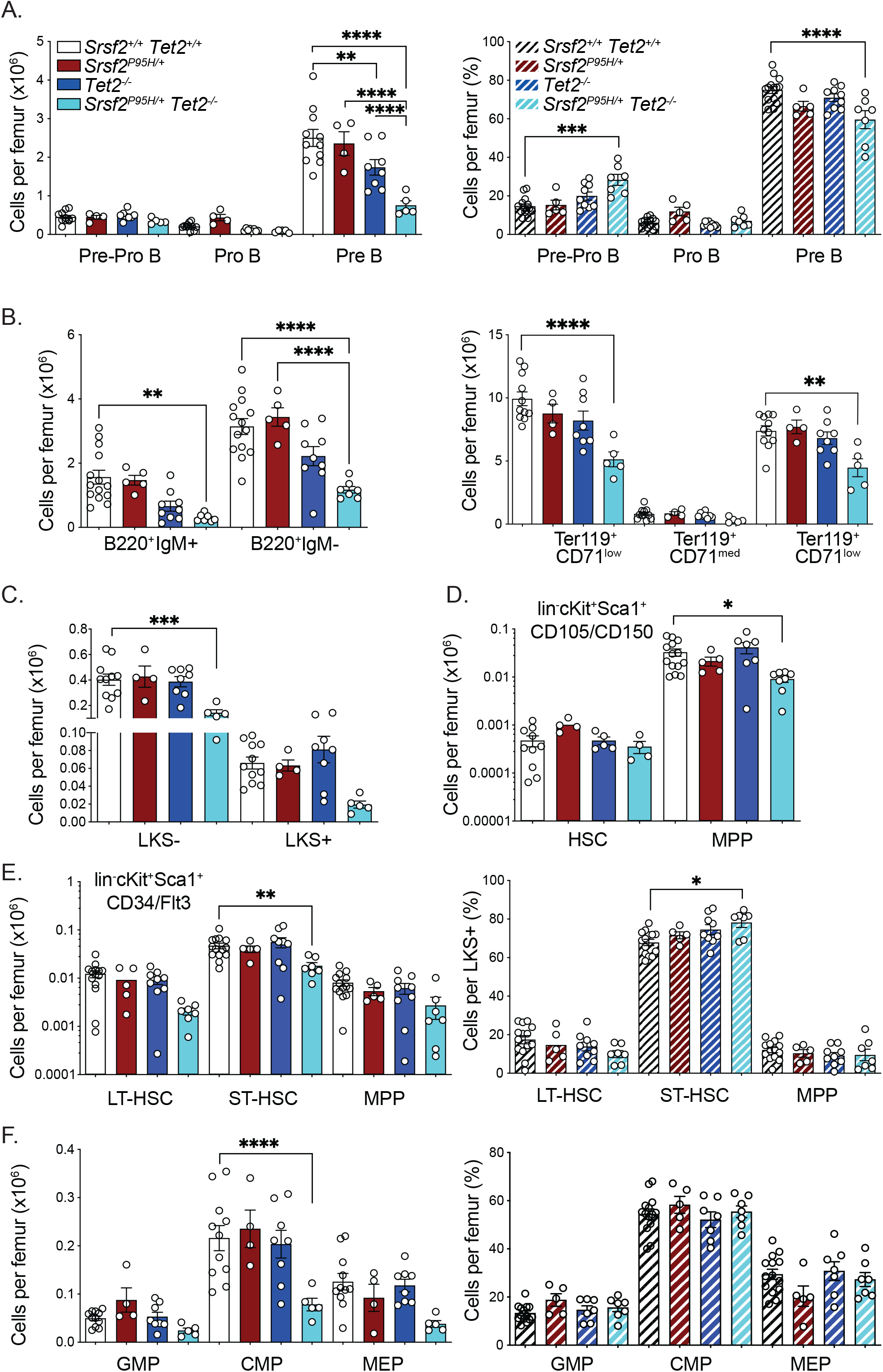
Concurrent mutation of *Srsf2*^*P95H/+*^ and deletion of *Tet2* leads to decreased Pre-B cells, decreased erythroid blasts and decreased stem and progenitor cell populations in the bone marrow. (**A**) Number and percentage of B progenitors in the bone marrow. (**B**) Number of B cells and erythroid blasts in the bone marrow. (**C**) Lin^-^c-Kit^+^Sca-1^-^ (LKS^-^) and Lin^-^c-Kit^+^Sca-1^+^ (LKS^+^) cells. (**D**) Number of hematopoietic stem cells (HSC) and multipotent progenitors (MPPs) using the lin^-^cKit^+^Sca1^+^CD105/CD150 phenotype (**E**) Number and percentage of long-term (LT-) and short-term hematopoietic stem cells (ST-HSC) and MPPs using the lin^-^cKit^+^Sca1^+^CD34/Flt3 phenotype. (**F**) Number and percentage of granulocyte macrophage progenitors (GMP), common myeloid progenitors (CMP) and myelo-erythroid progenitors (MEP). N=5-13 per genotype. IgM, immunoglobin M. Presented as mean +/-standard error of mean. One-way ANOVA performed against wild-type cells. \**P*<.05, \*\**P*<.01, \*\*\**P*<.001, \*\*\*\**P*<.0001.

Coincident with the reduction in BM cellularity in the *Srsf2/Tet2* compound mutant, there was a significantly reduced number of lineage^-^c-Kit^+^Sca-1^-^ (LKS-) cells (Figure 2C). Within the stem and progenitor containing lineage^-^c-Kit^+^Sca-1^+^ (LKS+) population, there was no apparent change in either absolute number or proportions of phenotypic long-term HSCs using multiple phenotyping approaches (Figure 2D-E). However, the percentage of phenotypic ST-HSCs was significantly increased in the double mutant mice (Figure 2E). Within the LKS-population, the double mutant mice had a significant reduction in the numbers of common myeloid progenitors (CMP), while GMP and MEP were not significantly altered (Figure 2F). Additionally, there were no alterations in other phenotypic myeloid and erythroid progenitor populations of the double mutant mice (Figure 2G). These analyses demonstrate that with aging in the setting of native hematopoiesis, *Srsf2*/*Tet2* double mutant cells develop a myeloid bias and expansion of the ST-HSCs.

In addition to the stem cell targeteing h*Scl-*CreER, we generated double mutant mice with broadly tissue tropic *Rosa26*-CreER^T2^. Whilst only a small cohort of mice has been analyzed at 62 weeks post mutation activation, the *Srsf2/Tet2* compound mutant mice had fewer B cells but no other significant changes in the PB. There was, however, a lower number of nucleated erythroid progenitors and Pro-B and Pre-B cells in the BM and an increased number of myeloid cells in the spleen (Supplemental Figure 2).

### Myeloid bias is a cell autonomous feature of *Srsf2*^*P95H/+*^ *Tet2*^*-/-*^ mutation

To determine the effect of stress on the *Srsf2*^*P95H/+*^ *Tet2* ^*Δ/Δ*^ compound mutant HSCs, BM transplantation was carried out. Both non-competitive and competitive transplants using whole BM of h*Scl-*CreER^T^ *Srsf2*^*P95H/+*^ *Tet2* ^*Δ/Δ*^ mice at 20 weeks after activation of the mutations were performed, along with transplant of tamoxifen-treated age-matched BM from Cre-positive wild-type (WT, +/+), single *Srsf2* mutant (*Srsf2*^*P95H/+*^) and single *Tet2* null (*Tet2* ^*Δ/Δ*^) mice. For non-competitive transplants, a total of 2 million whole BM cells (donor cells are CD45.2+) were intravenously injected into irradiated congenic recipient mice (CD45.1+) (Figure 3A). For competitive transplants, 1 million donor whole BM cells (CD45.2+) along with 1 million BM cells from a wild-type donor (CD45.1/CD45.2+; competitor cells; not age matched to mutant cell donor; 8-20 weeks of age) were intravenously injected into irradiated congenic recipient mice (CD45.1+) (Figure 3A).

**Figure 3.**
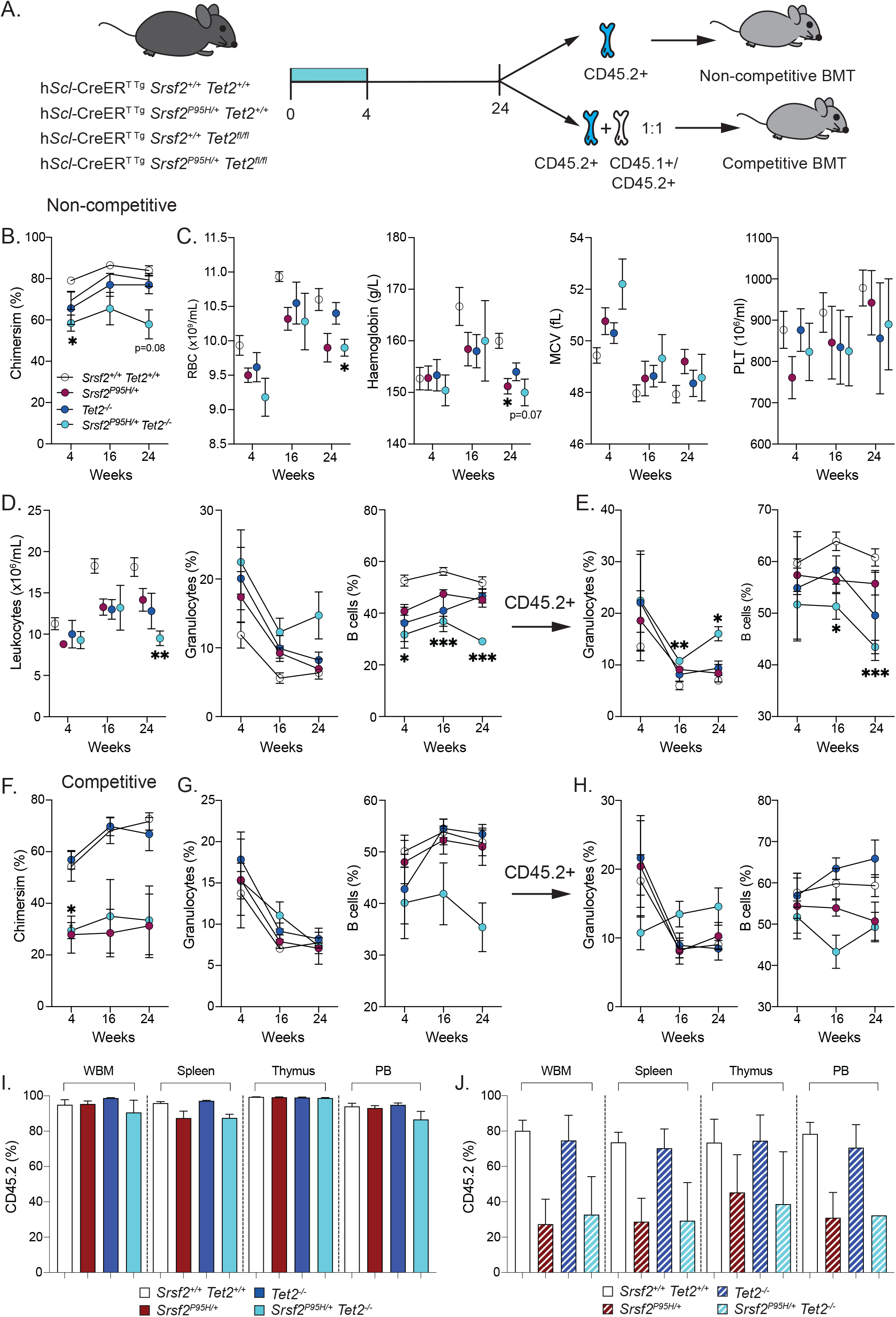
*Srsf2*^P95H/+^ *Tet2*^-/-^ cells have decreased engraftment capacity and transplantation leads to myeloid bias with B cell suppression in recipients. (**A**) Schematic illustration of non-competitive and competitive transplantation (B) Percent chimerism in non-competitive transplant. (**C**) PB indices at day 0 and 12- and 20-weeks post transplantation. (**D-E**) The total number of leukocytes and the percentage of granulocytes and B cells in total PB or CD45.2+ cells. (**F**) Engraftment of *Srsf2*^P95H/+^ *Tet2*^-/-^ cells in competitive transplantation. (**G-H**) The percentage of granulocytes and B cells in total leukocytes and CD45.2+ cells. (**I**) Engraftment of *Srsf2*^P95H/+^ *Tet2*^-/-^ cells across hematopoietic tissue at 1 year post non-competitive transplantation. (**J**) The percentage of donor *Srsf2*^P95H/+^ *Tet2*^-/-^ cells (CD45.2+) at 1 year post competitive transplantation. Number of recipients= 3-6 per genotype. Presented as mean +/-standard error of mean. One-way ANOVA performed against wild-type cells. \**P*<.05, \*\**P*<.01, \*\*\**P*<.001, \*\*\*\**P*<.0001.

Upon non-competitive transplantation, the *Srsf2*/*Tet2* compound mutant cells achieved a stable, but lower level of engraftment compared to WT, and this remained low at 24 weeks post-transplantation (Figure 3B). By 24 weeks post-transplant, the double mutant recipients had decreased leukocytes, RBCs and Hb in the PB, indicative of cytopenia and anemia (Figure 3C-D). Assessing lineage distribution, the double mutant BM recipients had significantly lower B cell frequencies throughout the analysis post-transplant. Within the CD45.2+ donor population, there was a significantly increased percentage of granulocytes and a significant decline in B cells at 16- and 24-weeks post transplantation (Figure 3E).

Upon competitive transplantation, in contrast, the double mutant cells had significantly reduced engraftment compared to WT and *Tet2* ^*Δ/Δ*^ BM recipients, with a level of reconstitution comparable to the *Srsf2*^*P95H/+*^ single mutant (Figure 3F). While the percentage of granulocytes was unchanged, the B cells frequency was lower in the double mutant recipients (Figure 3G). The granulocyte percentages within the CD45.2+ population were elevated, but not significantly (Figure 3H). At one year post transplantation, donor cells remained stably engrafted in all hematopoietic tissues in non-competitive recipients while the *Srsf2* single mutant and the compound mutant cells had a lower level of engraftment upon competitive transplant (Figure 3I-J).

To investigate the consequences of long-term mutation of *Srsf2/Tet2*, we performed non-competitive transplantation with whole BM from h*Scl-*CreER^T^ *Srsf2*^*P95H/+*^ *Tet2*^*-/-*^ mice at 56 weeks after gene mutation (Supplemental Figure 3A). In contrast to the 20-week donor marrow, the engraftment of aged *Srsf2*/*Tet2* compound mutant donor BM gradually declined and was significantly lower than the WT BM at 24 weeks post-transplant (Supplemental Figure 3B). Double mutant recipients had macrocytic RBCs and decreased leukocytes in the PB. Within the leukocyte fraction, the percentage of granulocytes was increased while B cells were decreased in the double mutant recipients (Supplemental Figure 3C-D). Within the CD45.2+ donor fraction, the double mutant BM had a significantly higher granulocytic than B cell output (Supplemental Figure 3E). Overall, the transplantation analysis demonstrated that myeloid bias is a cell-intrinsic consequence of *Srsf2*^*P95H/+*^ *Tet2*^*-/-*^ mutation, and that the *Srsf2*/*Tet2* compound mutant HSCs have a reduced competitive transplant capacity.

### *Srsf2^P95H/+^ Tet2^-/-^* leads to development of CMML *in vivo*

During aging, four mice developed phenotypes consistent with disease progression, one (#359) from an aging cohort of h*Scl-*CreER^T^ *Srsf2*^*P95H/+*^ *Tet2* ^*Δ/Δ*^ and, two (#836, 835) from the non-competitive transplantation of two pooled BM from 20 weeks post tamoxifen treated h*Scl-*CreER^T^ *Srsf2*^*P95H/+*^ *Tet2* ^*Δ/Δ*^ mice. The last case (#290) was from non-competitive BM transplantation from a 56-week post-tamoxifen treated h*Scl-*CreER^T^ *Srsf2*^*P95H/+*^ *Tet2* ^*Δ/Δ*^ mouse. Of the four mice that developed advanced disease, #359 became moribund at 52 weeks post tamoxifen treatment, while #835 and #836 were identified at 52 weeks and 12 weeks post-transplantation, respectively. Mouse #290 became moribund at 28 weeks post-transplantation (Figure 4A). At the time of analysis, #835 and #836 had significantly increased leukocyte counts while #359 and #290 had normal leukocyte numbers in the PB. The BM cellularity of the four mice was normal or lower than the age-matched WT context specific control (for #359 the average BM cellularity for control animals was 32.9±1.2 × 10^6^; for #835 and #836 the average BM cellularity for transplant controls was 27.8±1.4 × 10^6^, for #290 the average BM cellularity for transplant controls was 38±18 × 10^6^) (Figure 4B). Both #359 and #835 displayed splenomegaly and increased splenic cellularity (Figure 4B-C). Mouse #836 had increased splenic size (183mg), whilst #290 had a subtle increase in spleen weight and cellularity.

**Figure 4.**
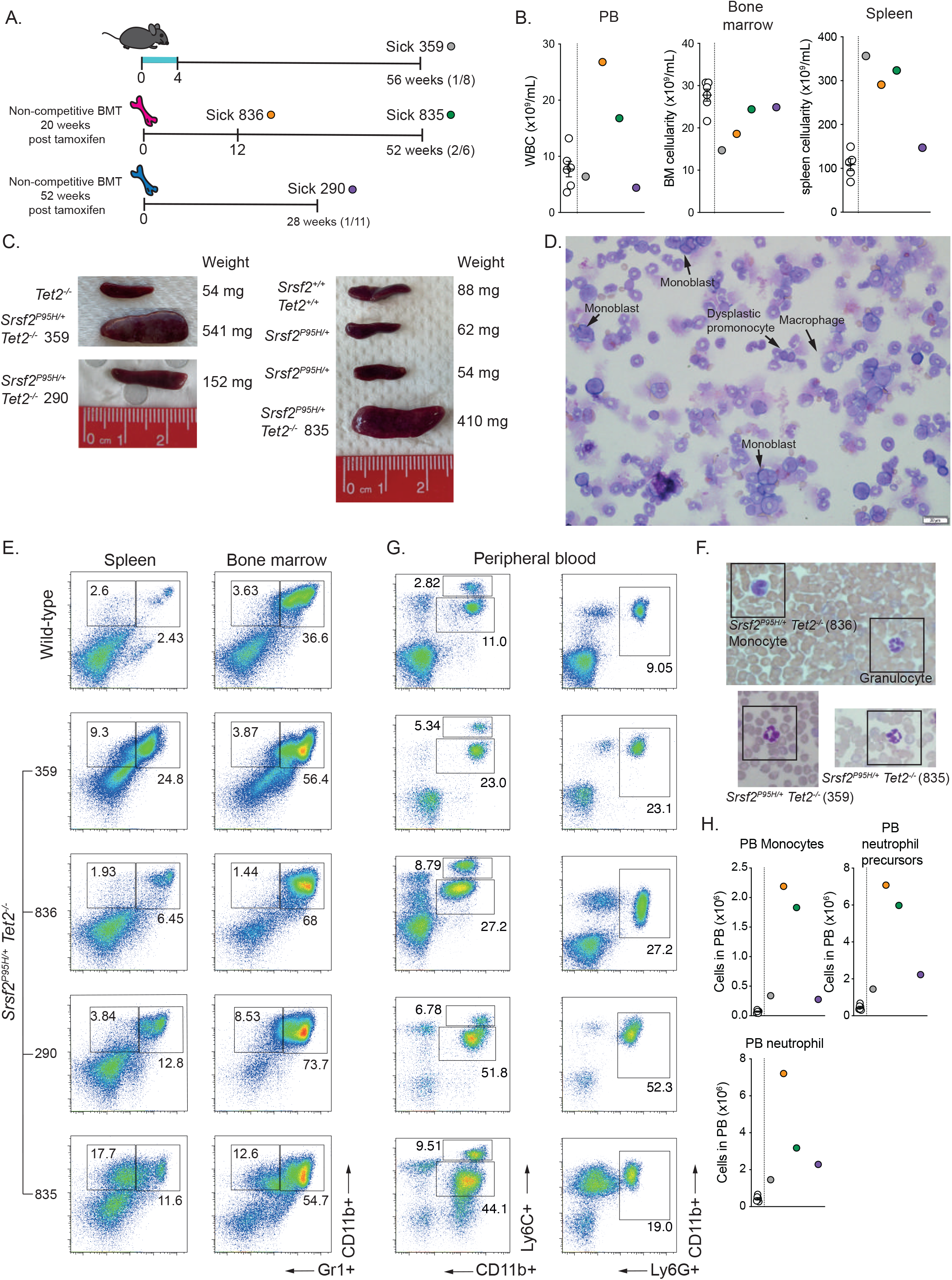
Sick mice display features consistent with CMML including splenomegaly, dysplastic neutrophils and increased numbers of immature monocytes. (**A**) A timeline of disease progression post mutation initiation or post transplantation. In brackets the number of diseased animals vs the number of aged animals for each experimental condition. (**B**) Cellularity of peripheral blood, bone marrow and spleen of moribund mice. (**C**) Images of spleen of primary (#359) and transplant aged (#835) mice compared to other genotypes in the cohort. (**D**) Bone marrow cytospin of primary (#359) at the time of analysis. Cells from the monocytic lineage at different stages of maturation. May-Grünwald-Giemsa staining. Image taken at 20x. (**E**) FACS plots of BM and spleen of moribund mice showing myeloid expansion. (**F**) Representative images of dysplastic neutrophils in the peripheral blood of moribund mice. May-Grünwald-Giemsa staining. Image taken at 40x. (**G-H**) FACS plots and quantification of monocytes and neutrophil precursors in the peripheral blood of sick mice compared to age matched wild-type.

Examination of the BM cytospins of the primary non-transplanted *Srsf2*^*P95H/+*^ *Tet2* ^*Δ/Δ*^ mouse (#359) showed an increased percentage of pro-monocytes and monoblasts (10-15%) along with occasional dysplastic monocytes (Figure 4D). There were elevated numbers of myeloid cells present in the BM and spleens of all four animals (Figure 4E) PB smears demonstrated that there were dysplastic (hypersegmented) neutrophils present in samples from #359, 835 and 836. For transplant recipients #835 and #836, eosinophils, promonocytes and monoblasts (#836) were identified in the PB smears (Figure 4F, Supplemental Figure 4A, 4C). Besides morphological analysis, immunophenotypic analysis of the PB, BM and spleen showed that there was a marked increase in granulocytes, monocytes (Ly6C^hi^, CD11b^+^) and neutrophil precursors/eosoinophils (Ly6C^med^, CD11b^+^) across all tissues (Figure 4F-H). Diseased mice presented with a reduced expression of CD45RB, evidence of mis-spliced CD45 protein resulting from SRSF2^P95H^ mutation^17,30^ (Supplemental Figure 4B). We confirmed the disease contributing cells are *Srsf2/Tet2* mutant cells as they were CD45.2+ positive at the time of analysis (Supplemental Figure 4D). These collective phenotypes are consistent with the pathological features of CMML, most closely aligning to CMML-2 (5-19% peripheral blasts and 10-19% marrow blasts, WHO diagnostic criteria)^31^.

### Exome sequencing reveals additional mutations linked to human CMML

A subset of *Srsf2/Tet2* compound mutant animals developed a CMML like disease phenotype. We then sought to determine if this resulted from additional mutations being acquired during disease progression. Exome sequencing of whole BM samples from two primary *Srsf2/Tet2* mutant samples with a myeloid bias phenotype, but without evidence of frank disease, and the four CMML samples described previously was undertaken. We compared them to either age-matched WT or a non-transplanted mutant sample (#290 vs #356) (Figure 5A, Supplemental Table 1). Genotyping confirmed the mutation of *Srsf2*^*P95H/+*^ and between 50-90% recombination and deletion of *Tet2* in the BM samples used in this analysis (Figure 5B). Interestingly, the mice that demonstrated disease progression also had the most complete recombination of *Tet2* (Figure 5B and Supplemental Dataset 1), suggesting that the variability in *Tet2* deletion may contribute to the low penetrance of disease in the compound mutant mice. From sequencing, the majority of mutations detected were missense predicated to have moderate to high impact ^29^ (Figure 5C). A range of mutations in genes reported in human myeloid malignancy are seen, with a trend to more mutations amongst the samples with evidence of CMML compared to those with myeloid bias but not frank disease phenotypes (Figure 5C) ^32^. The mutations cover a range of pathways and individual genes known to be mutated in human CMML, however only a small number of samples have been assessed. Collectively, the results demonstrate that the combined mutations of *Srsf2* and *Tet2* in HSCs is able to initiate a disease, in a subset of animals, that has phenotypic and genetic similarity to human CMML.

**Figure 5.**
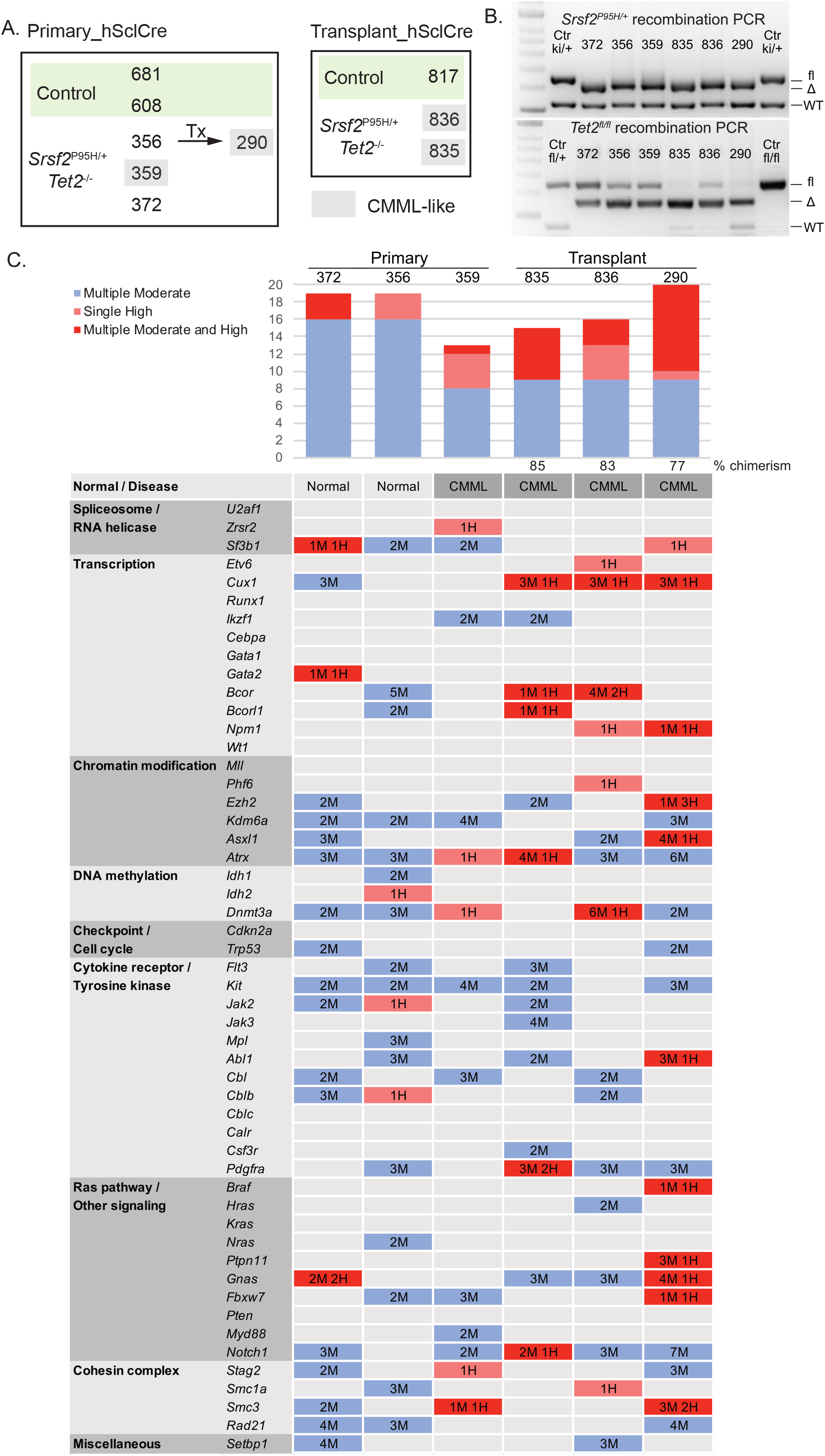
Exome sequencing of CMML mice reveals additional mutations acquired during disease development. (**A**) Samples of CMML-like mice and respective control that were used in exome sequencing. (**B**) PCR showing recombination of *Srsf2*^P95H/+^ and *Tet2*^-/-^ allele in the bone marrow of analyzed mice. (**C)** Somatic variant detection in a panel of 52 genes frequently mutated in myeloid cancers. “Tumor” samples were compared to matched “Normal”. The number of missense mutations with Moderate Impact (M, blue shade), single High Impact (H, light red shade), or Multiple Moderate and High Impact (M/H, dark red shade) are indicated for each gene.

## Discussion

Herein we describe the generation and characterization of a *Srsf2 Tet2* mutant murine model which reproduces features of CMML. Aging of the compound mutant mice resulted in a more overt myeloid bias and B cell suppression compared to either *Srsf2* mutation or *Tet2* loss alone. Proceeding disease development, we observed phenotypes unique to the compound mutants including hypocellular BM and monocytosis. In the reported single mutant models, none of the *Tet2* models had decreased bone marrow cellularity ^15,16,33,34^, while our *Srsf2* model showed a trend towards lower BM counts in the *Srsf2*^*P95H/+*^ mice ^17^. In the *Srsf2 Tet2* double mutants, reduced progenitor, lymphoid and erythroid cell numbers are likely to be the source of hypocellularity. Both *SRSF2* and *TET2* mutations are associated with monocytosis in patients^5,32,35,36^. The compound mutant mice displayed monocytosis, as well as increased immature pro-monocytes and monoblasts, suggesting a synergistic effect of *Srsf2 Tet2* mutation on the early expansion of the monocytic blast population. Similar to the primary mice, recipients of *Srsf2*^*P95H/+*^*Tet2* ^*Δ/Δ*^ mutant BM developed marked myeloid bias and B cell suppression, evidence that the lineage bias was cell-autonomous. Compound mutant HSCs were able to achieve stable long-term engraftment in a non-competitive setting. Surprising, compound mutant BM showed a disadvantage when competitively transplanted with wild-type BM, most similar to the *Srsf2* single mutant cells. Upon aging and or transplant, we observed that a subset of animals developed features consistent with CMML. The disease phenotypes and additional mutations identified through exome sequencing confirmed the establishment of a murine model with core features of human CMML.

We observed a long latency and incomplete penetrance of CMML, with the earliest disease becoming apparent one year after gene mutations. Most double mutant animals exhibited myeloid expansion with one out of eight aged to 52-60 weeks non-transplant mice (14%), two out of six (33%) of non-competitive transplant recipients and one out of eleven (9%) aged-donor-BM transplant recipient showed evidence of progression to CMML. The long latency, including the apparent increased penetrance following transplant, indicates the necessity to acquire additional mutations that facilitate the evolution and out-growth of clones with disease-initiating potential. In humans, both *SRSF2* and *TET2* mutations are reported in clonal hematopoiesis of intermediate potential (CHIP) and even in some healthy individuals ^4,37-39^. In CMML, both mutations are considered as the ancestral mutations that occur very early in disease initiation. The present data using timed somatic mutation and longitudinal analysis in the mouse demonstrate that the effects of either SRSF2^P95H^ or TET2 loss alone are, overall, well tolerated and whilst myeloid biasing do not result in profound disruptions to hematopoiesis. SRSF2^P95H/+^ compromises competitive transplant potential of HSCs but TET2 loss does not. The HSC transplant potential of the compound mutant cells were most similar to the *Srsf2*^*P95H/+*^ single mutant, demonstrating the loss of TET2 is not sufficient to rescue the reduced competitiveness of *Srsf2*^*P95H/+*^ HSCs.

The most distinctive feature of compound SRSF2 and TET2 mutation was monocytosis and monocytic expansion. The data demonstrate that the combined mutation of SRSF2 and loss of TET2 results in a myeloid shift and a fertile cellular environment for acquisition of additional mutations and disease transformation ^3^. Recent evidence demonstrates that along a spectrum from clonal hematopoiesis of indeterminant potential (CHIP) to CMML the median number of mutations increases. CMML has a median of 4 consequential mutations (range, 1-6 mutations)^39^. The murine data indicate that while SRSF2 and TET2 mutations can co-operate to drive development of a disease with key characteristics of CMML, the penetrance and latency of the model could be further improved by inclusion of mutations in *ASXL1* and *RAS* pathway components as seen in human. A recently reported model demonstrated higher penetrance of a CMML like disease by combining an *Asxl1* mutation with the G12D mutation in *Nras*, a gain of function mutation that confers a significant proliferative advantage ^19^. We would predict that an increased penetrance and decreased latency of disease would be elicited in the *Srsf2/Tet2* compound mutant model by introducing a mutation conferring a proliferative advantage. Consistent with this concept, in preliminary data from a small cohort of animals, we overserved a more profound and rapid myelomonocytic expansion when the *Srsf2*^*P95H/+*^ mutation was couple with a *Cbl* null allele (Supplemental Figure 5).

One intriguing finding was that when assessing the *Srsf2* and *Tet2* recombination efficiency, we noted that the *Tet2* allele often did not achieve full recombination following tamoxifen treatment. In contrast, the *Srsf2*^*P95H/+*^ allele was efficiently recombined by 4 weeks of tamoxifen treatment. Consistent with our observations, in the *Tet2* models reported by Quivoron et al., and Izzo et al., there was also evidence of incomplete recombination of *Tet2* ^16,40^. These studies used different Cre drivers and means to elicit gene deletion (e.g. *Mx1*-Cre and pIpC compared to h*Scl-*CreER and tamoxifen), yet a common finding is the incomplete recombination of the *Tet2* floxed allele. This appears specific to the recombination efficiency at the *Tet2* floxed allele within this mouse line. Remarkably, there seems to be no selective pressure for selection for fully TET2 null cells or for loss-of-heterozygosity of the *Tet2* allele, even in the presence of co-operating mutations. As TET2 is expected to be a loss-of-function mutation in human disease, whether this partial loss of TET2 activity in our model could contribute to the long latency is at present unknown^41^.

By combining the *Srsf2*^*P95H/+*^ mutant and *Tet2* loss of function alleles, we demonstrate *in vivo* co-operativity of *Srsf2*^*P95H/+*^ and *Tet2* ^*Δ/Δ*^ in native hematopoiesis and synergized phenotypes, such as monocytosis and BM hypocellularity. A subset of animals progress to develop a disease unique to the compound mutants with features consistent with CMML. The rate of disease progression was higher following transplantation, potentially allowing expansion of pre-existing small clones harboring multiple mutations from the primary donors. In conclusion, we describe a new pre-clinical model that recapitulates the genetic association of *SRSF2* and *TET2* mutations seen in patients and mirrors aspects of human CMML. This is a significant first step toward building high fidelity autochthonous models to understand the evolution from clonal hematopoiesis to CMML development and to test new therapeutic agents.

## Acknowledgements

The authors would like to thank L. Purton for discussion and comments; St. Vincent’s Hospital BioResources Centre for care of experimental animals; St. Vincent’s Institute Flow Cytometry Core Facility. This work was supported by the Cancer Council of Victoria (C.R.W., A.M.C.; APP1126010), Victorian Cancer Agency Research Fellowship (C.R.W.; MCRF15015); and in part by the Victorian State Government Operational Infrastructure Support (OIS) to St Vincent’s Institute.

## Authorship contribution

J.J.X, M.F.S, C.R.W conceptualized the study; J.J.X, W.Y.L., M.F.S and C.R.W designed the experiments; J.J.X, A.M.C, M.W, M.F.S and C.R.W performed the experiments.; J.J.X and A.M.C produced the figures; J.J.X wrote the original manuscript; J.J.X, A.M.C, M.W, W.Y.L., M.F.S and C.R.W reviewed and edited the manuscript; A.M.C and C.R.W were responsible for funding acquisition; M.F.S and C.R.W provided supervision.

