## Supplementary material for "*Srsf2^P95H/+^* co-operates with loss of TET2 to promote myeloid bias and initiate a chronic myelomonocytic leukemia like disease in mice": Supp Methods and Figures

**Contents:**

Page 3-5: Supplemental Methods and References
Page 6: Description of Supplemental Datasets S1
Page 7 - :  Supplemental Figures and legends S1-S5

### Methods

#### Ethics Statement

All animal experiments were approved by the AEC (AEC#001/16 and 007/19); St. Vincent's Hospital, Melbourne).

#### Animal Models

*Srsf2*$^{P95H/+}$ mice (C57BL/6NTac-*Srsf2*$^{tm2874(P95H)Arte}$) have been previously described and characterized[1-3].

Genotyping was performed by PCR as follows:

5'loxP: 6034_38: 5'-GTTATGATCCACACCTCTCACC-3',

6034_39: 5'-ATAAACGTTTATGTCCGCTACC-3';

To yield products of 327bp for WT, 446bp for an intact 5'loxP site, and 400bp for a recombined product.

3'loxP: 2240_31 hGHpA 5´2: 5-ACGTCCAGACACAGCATAGG-3',

6035_41: 5'-GGCTTGCAATTTGAGTATGG-3',

1260_1: 5'-GAGACTCTGGCTACTCATCC-3',

1260_2: 5'-CCTTCAGCAAGAGCTGGGGAC-3';

To yield products of 364bp (KI) for an intact 3'loxP site and 585bp (internal control PCR product using 1260_1 and 1260_2).

For sequence verification of P95H mutation:

6036_44: 5'-GCGCGTCTTCGAGAAATACG-3',

6036_43: 5'-ACGTGAACGAAGCGACAGG-3';

To yield products of 630bp (WT) and 630bp (PM) for a product that requires Sanger sequencing.

The introduced nucleotide change generates a PM unique Hpy188III restriction site which when digested yields a 363/184/46/37bp digest product (PM) or 547/46/37bp digest product (WT) from the PCR product used for sequencing (primers 6036_44 and 6036_43).

*Tet2*$^{fl/fl}$ mice were purchased from the Jackson Laboratory and genotyped as described on the Jackson Laboratory website (B6;129S-*Tet2*$^{tm1.1Iaai/J}$; strain #017573, Jackson Laboratory).

Recombination/deletion of exon 3 was verified by PCR using the primers:

13495: 5′-AAGAATTGCTACAGGCCTGC-3',

13496: 5′-TTCTTTAGCCCTTGCTGAGC-3',

Tet2 recombine_1: 5′-CCAATGACAGGCCCAAATTGT-3';

To yield products of 249bp for WT, 427bp for an intact 5'loxP site, and 350bp for a recombined product.

*c-Cbl*$^{-/-}$ (MGI:2180578) were provided by Wallace Langdon (UWA) and have been previously described[4].

Genotyping was performed by PCR as follows:

MM1 (common forward primer) - 5'TAGGCGAAACCTGACCAAAT3'

c-Cbl WT R1 (reverse primer for WT allele) - 5'CTC GCT CAG TCC CTG CTT AC3'

MM3 (reverse primer for KO allele) - 5'TGCTACTTCCATTTGTCACG3'

To yield products of: WT allele=456bp; KO allele= 288bp

h*Scl*-CreER$^{T}$, R26eYFP (strain #006148, Jackson Laboratory; [1,5-7]), and *Rosa26*-CreER$^{T2}$ (strain #008463, Jackson Laboratory; [8-10]) mice have been previously described and were genotyped as described on the Jackson Laboratory strain information datasets.

Congenic B6.SJL-Ptprc$^{aPep3b/BoyJArc}$ were purchased from the Animal Resources Centre (Canning Vale, Western Australia). Hetrozygous CD45.1/CD45.2 C57BL/6 mice were bred at St Vincent's Hospital, Melbourne. Tamoxifen containing food was prepared containing 400mg/kg tamoxifen citrate (Sigma or Selleckchem) with 5% sugar (refined white sugar) in irradiated standard mouse chow (Specialty Feeds, Western Australia). Tamoxifen containing chow was fed *ad libitum* during the treatment period then animals were returned to normal diet [1,9,10]. All lines were on a C57BL/6 background.

**Flow cytometry analysis and fluorescent activated cell sorting**

Peripheral blood was analyzed on a hematological analyzer (Sysmex KX-21N, Roche Diagnostics). Bones were flushed, spleens and thymi crushed and single cell suspensions were prepared[1,9]. Antibodies against murine Ter119, CD71, B220, IgM, Mac-1, Gr1, F4/80, CD43, CD19, CD4, CD8, CD25, CD44, Sca-1, c-Kit, CD34, FLT3, Fc$\gamma$R (CD16/32), CD41, Ly6C, Ly6G either biotinylated or conjugated with FITC, phycoerythrin, phycoerythrin-eFluor610, peridinin chlorophyll protein-Cy5.5, phycoerythrin-Cy7, allophycocyanin, allophycocyanin-eFluor780, eF660 or eF450 were all obtained from eBioscience. CD105 and CD150 were from BioLegend. CD45RB from BD Biosciences. Biotinylated antibodies were detected with streptavidin conjugated with Brilliant Violet-605 (Biolegend). Cells were analyzed on a BD LSRIIFortessa (BD Biosciences) and where applicable cells were sorted with either BD FACSAria or BDInflux (BD Biosciences). Results were analyzed with FlowJo software Version 10.0 (Treestar).

**Hematopathology**

May-Gunwald Giemsa or hematoxylin and eosin stained respectively peripheral blood films and bone marrow (both sections and cytospins) were assessed by a hemato-pathologist.

**Supplemental Datasets:**

**Dataset S1:** Summary of exome capture data and mutations identified (related to Figure 5).

Xu et al. Supp figure 1

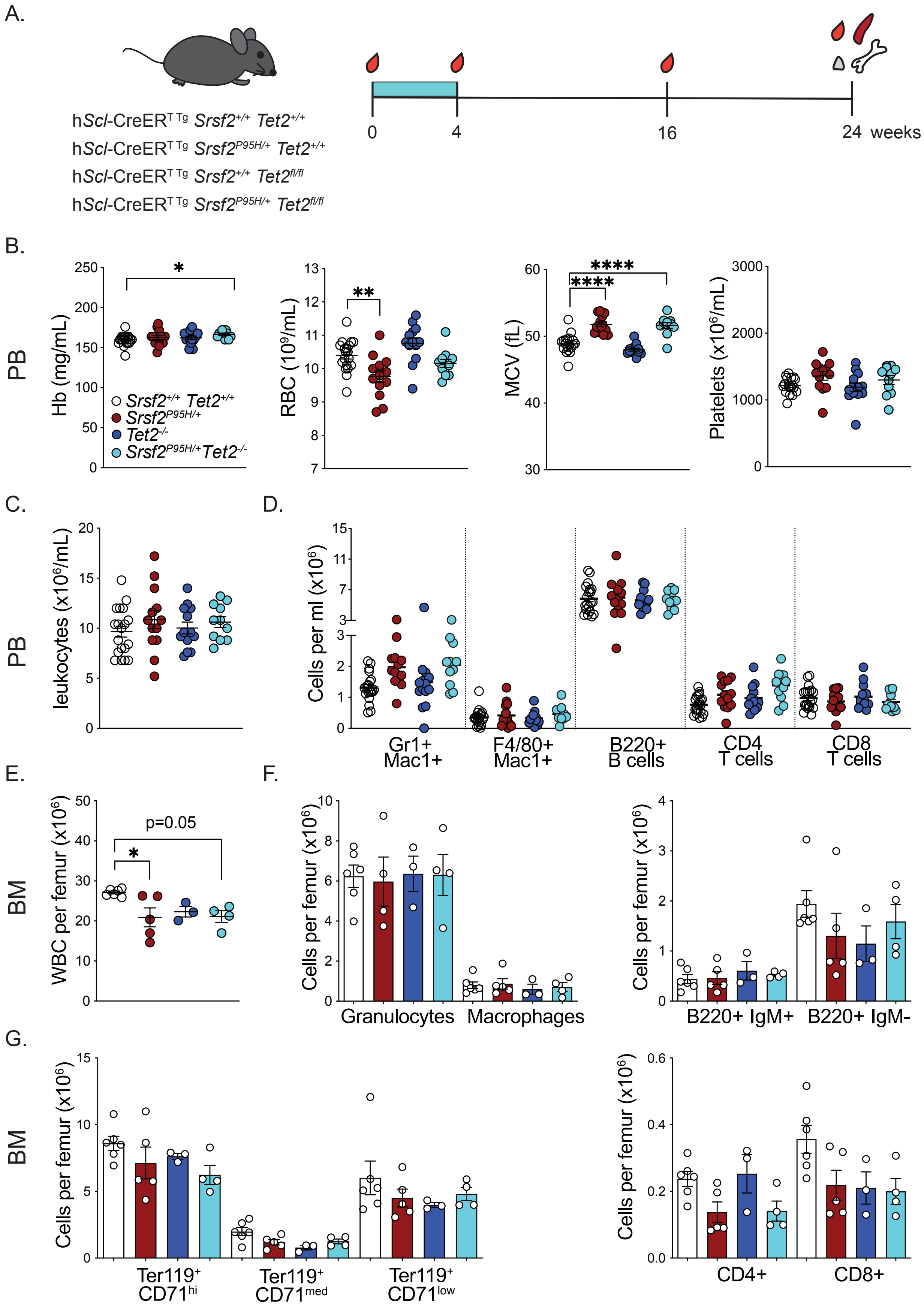

**Supplemental Figure 1. Concurrent mutation of *Srsf2*$^{P95H/+}$ and deletion of *Tet2* leads to macrocytic changes after short-term mutation activation**. (**A**) Schematic of h*Scl*-CreER *Srsf2*$^{P95H/+}$ *Tet2*$^{-/-}$ experiments. (**B**) PB indices after 20 weeks of Cre activation. PB leukocyte counts (**C**) and lineage distribution (**D**) after 20 weeks of Cre activation (n≥11 per genotype). (**E**) Bone marrow cellularity per femur. (**F**) Number of myeloid and B lymphoid cells per femur. (**G**) Number of erythroid precursors and T cells per femur (n≥3 per genotype). IgM, immunoglobin M. Presented as mean +/- standard error of mean. One-way ANOVA performed against wild-type cells. *$P$<.05, ****$P$<.0001.

Xu et al. Supp figure 2

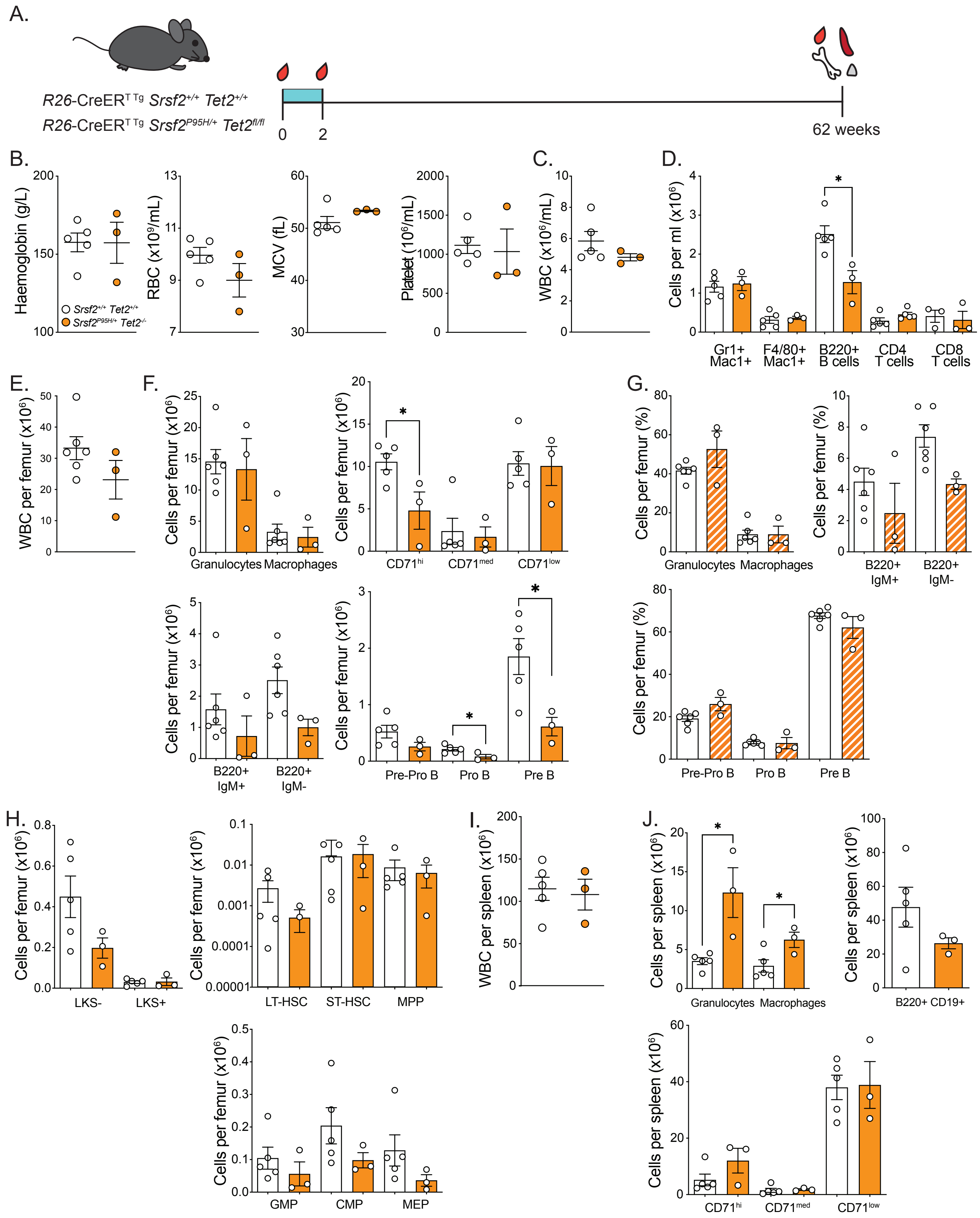

**Supplemental Figure 2. Widespread co-expression of *Srsf2*$^{P95H/+}$ and *Tet2*$^{-/-}$ leads to decreased B cell and erythroid blast populations**. (**A**) Schematic outline of *R26*-CreER$^{T2}$ $^{Tg}$ *Srsf2*$^{P95H/+}$ *Tet2*$^{-/-}$ experiments. PB indices of *Srsf2*$^{P95H/+}$ *Tet2*$^{-/-}$ mice (**B**), leukocyte counts (**C**) and lineage counts (**D**) compared to age-matched controls after one year of Cre activation. (**E**) Bone marrow cellularity after one year of mutation activation. The number of myeloid, B lymphoid, erythroid cells (**F**) and their percentages (**G**) in the bone marrow. (**H**) The number of stem and progenitor populations in the bone marrow. (**I**) The spleen cellularity after one year of mutation activation. (**J**) The number of myeloid, B lymphoid lineages and erythroid blasts in the spleen. Presented as mean +/- standard error of mean. Student t-test. *$P$<.05.

Xu et al. Supp figure 3

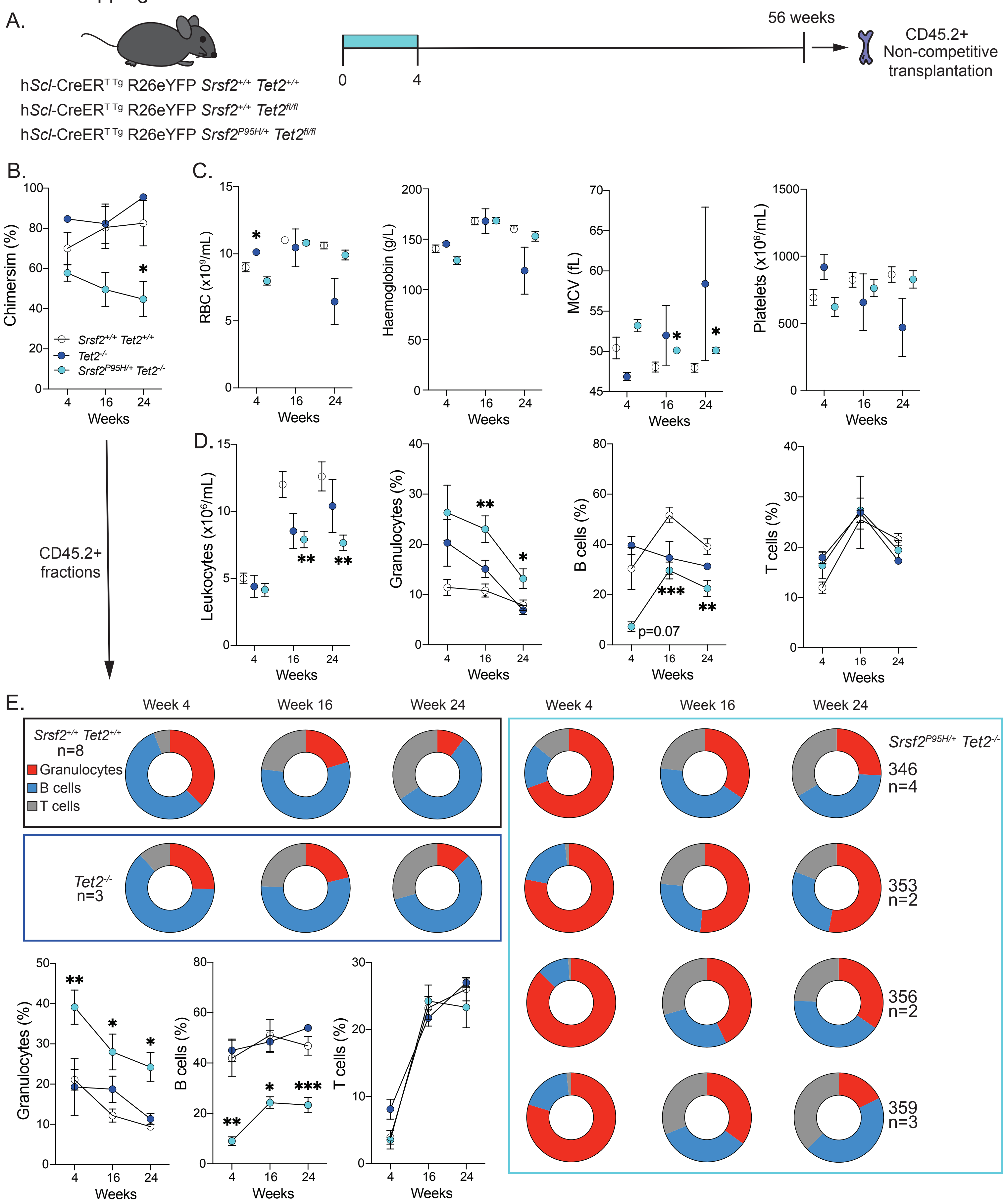

**Supplemental Figure 3. Aged *Srsf2*$^{P95H/+}$ *Tet2*$^{-/-}$ cells have decreased engraftment capacity and transplantation leads to myeloid bias in recipients**. (**A**) Schematic illustration of non-competitive transplantation experiments with aged bone marrow (52 weeks post mutation activation). (**B**) Aged *Srsf2*$^{P95H/+}$ *Tet2*$^{-/-}$ cells achieved a lower engraftment in non-competitive transplantation. (**C**) PB indices of recipients up to 20 weeks post transplantation. (**D**) PB leukocyte counts and lineage distribution of non-competitive transplant recipients. (**E**) PB lineage distribution of donor (CD45.2+) population in non-competitive transplant recipients. Number of recipients = 2-8 per genotype. WT recipients received BM from two individual wild-type donors (n=5 for 681 and n=3 for 281 and results were pooled). Other genotype recipients received BM from a single donor. Presented as mean +/- standard error of mean. One-way ANOVA performed against wild-type cells. *$P$<.05, **$P$<.01, ***$P$<.001, ****$P$<.0001.

Xu et al. Supp figure 4

A.

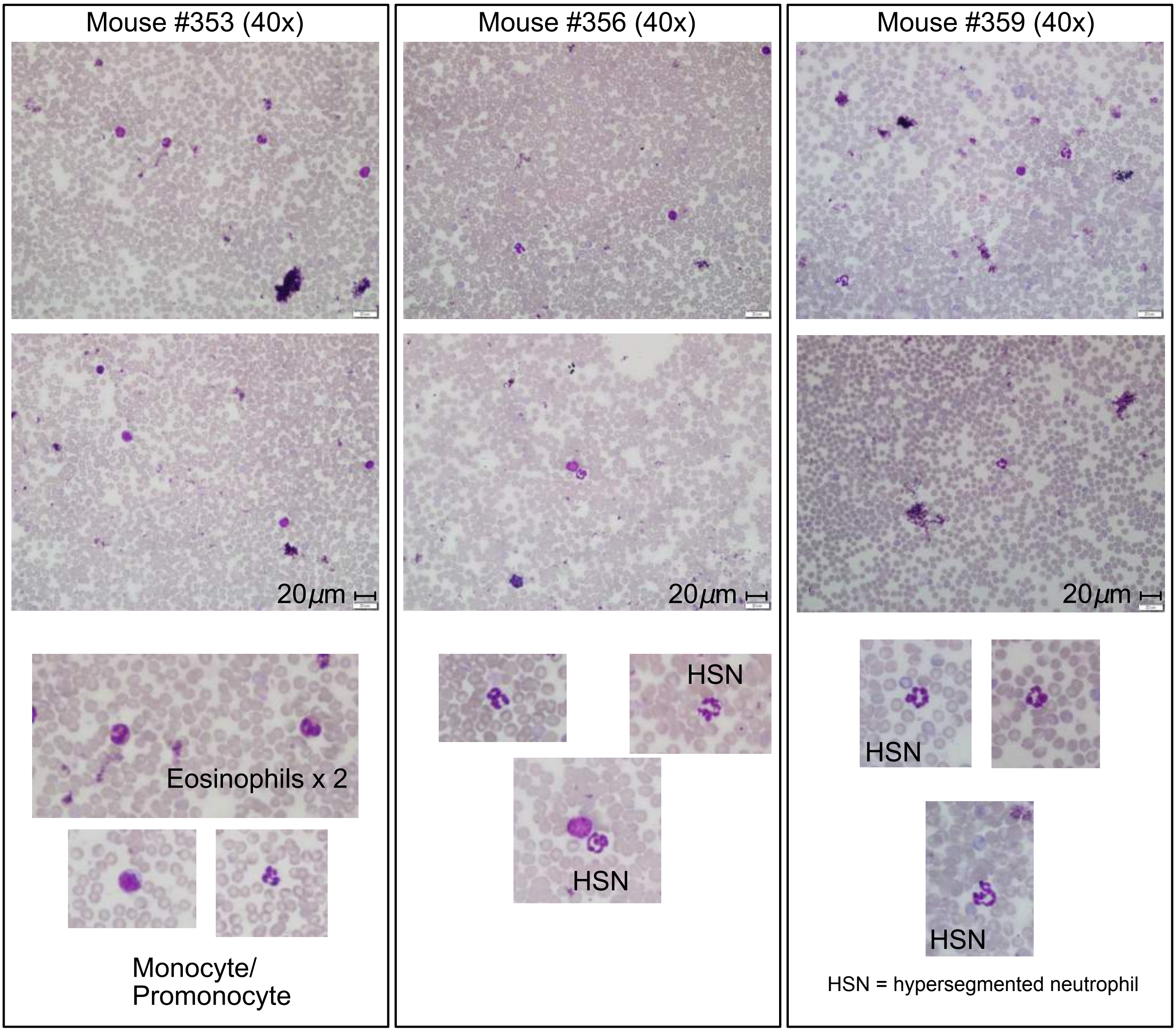


B.

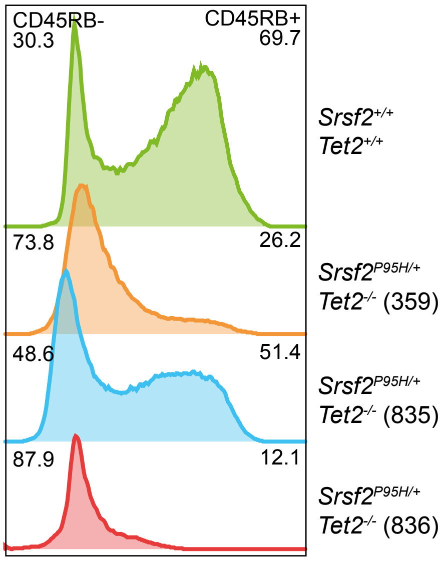

Xu et al. Supp figure 4 cont.

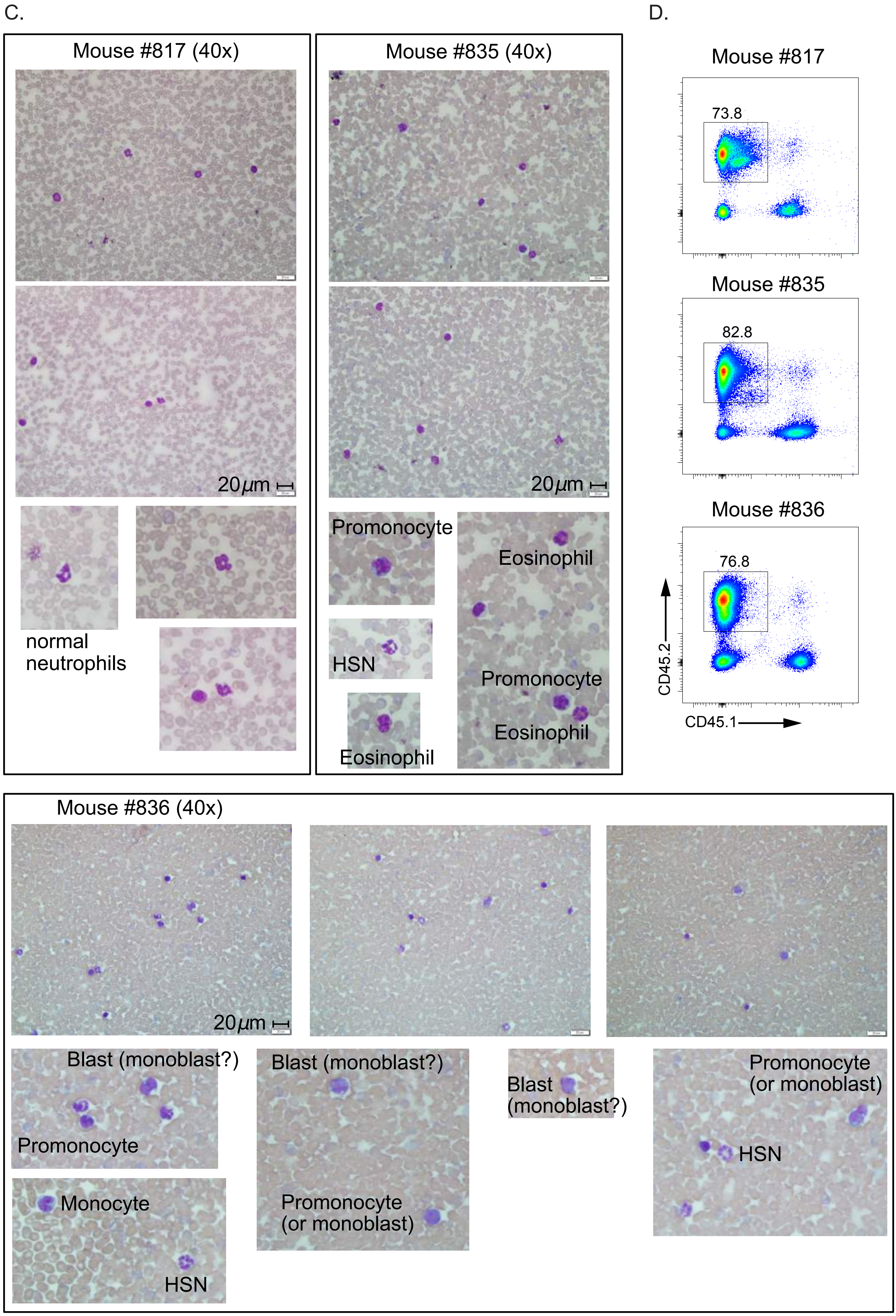

**Supplemental Figure 4.** Hematopathology of mice assessed by exome capture. (**A**) Blood film analysis of aged *Srsf2*$^{P95H/+}$ *Tet2*$^{-/-}$ mouse indicates presence of promonocytes, monocytes, eosinophiles and hypersegmented neutrophil (HSN). (B) Histogram of CD45RB isoform expression on splenocytes of the indicated mice. (C) Blood film analysis of transplant recipients. (D) Chimerism of the analyzed recipients at time of collection.

Xu et al., Supp figure 5

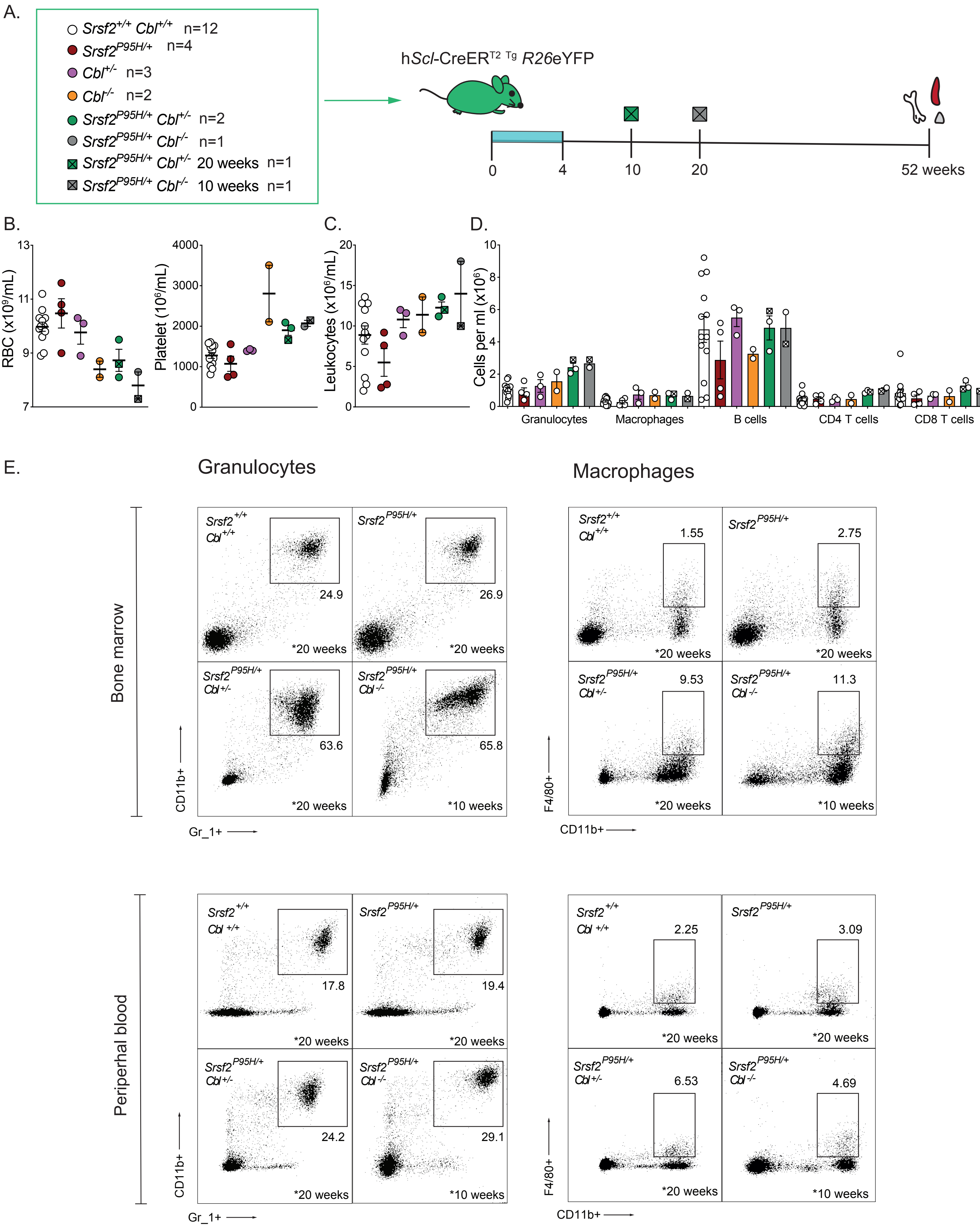

**Supplemental Figure 5. Co-expression of *Srsf2*$^{P95H/+}$ and *Cbl* deletion accelerates the development of myeloid-hyperplasia**. (**A**) Schematic illustration of h*Scl*-CreER *Srsf2*$^{P95H/+}$ *Cbl*$^{-/-}$ experiments. PB indices of *Srsf2*$^{P95H/+}$ *Cbl*$^{-/-}$ mice (**B**), leukocyte counts (**C**), and lineage output (**D**), compared to aged-matched Cre control. (**E**) FACS plots of granulocytes and macrophages of CMML-like *Srsf2*$^{P95H/+}$ *Cbl*$^{-/-}$ mice collected at 10-week or 20-week post mutation activation (n=1 per timepoint)
